# Evolution and adaptations of the seminal proteome in an insect with traumatic insemination

**DOI:** 10.64898/2026.04.15.718635

**Authors:** Martin D. Garlovsky, Oliver Otti, Klaus Reinhardt, Timothy L. Karr

## Abstract

The protein composition of sperm and seminal fluid are key to male fitness. However, we currently lack an understanding of the factors that shape seminal proteome composition. The common bedbug (*Cimex lectularius*) mates by traumatic insemination, subjecting the ejaculate to a unique selective environment as sperm traverse the female genital and paragenital system. We provide the first high-throughput proteomic characterisation of the sperm and seminal fluid proteome in a hemimetabolous insect and the first in-depth proteomic characterisation of the male bedbug reproductive system. Our analysis revealed conserved and unique features of the sperm and seminal fluid proteome with possible links to features of sperm behaviour linked to traumatic insemination. The sperm proteome showed elevated rates of molecular evolution, unlike most other studied species. Conversely, the sperm proteome also contained many conserved proteins. Notably, we found an expansion of Sperm-leucylaminopeptidases (S-Laps) in bedbugs and other hemimetabolous insects, suggesting the origin of S-Laps is perhaps even more ancient than previously thought. Using *in silico* protein-ligand binding predictions, we show that S-Laps have likely retained catalytic activity. Our results provide a list of candidate proteins involved in reproduction and a foundation for future studies of this expanding global pest.

## INTRODUCTION

The proteomic composition of sperm and seminal fluid is central to fertility and fitness (Snook 2005; Perry et al. 2013). Proteins are a major component of gametes, in large part dictating spermatozoa form and fertilisation capacity (Dorus and Karr 2009). Likewise, seminal fluid proteins transferred along with sperm in the ejaculate aid sperm transport and storage and affect female physiology and morphology after mating (Wigby et al. 2020). However, until recently the tools needed to study the molecular composition of the ejaculate were lacking (Karr 2019). Consequently, our understanding of the factors shaping seminal proteome composition and evolution remain poorly understood.

High-throughput shotgun proteomics studies have revealed the sperm and seminal fluid proteome contain hundreds or thousands of proteins (Wigby et al. 2020; Pini et al. 2025). Recent studies have highlighted a nuanced landscape of the seminal proteome. For instance, some seminal fluid proteins may have a testis origin (Garlovsky et al. 2022; McCullough et al. 2022). Many reproductive proteins show elevated rates of molecular evolution (Swanson and Vacquier 2002; Dapper and Wade 2020; Patlar et al. 2021). However, the sperm and seminal fluid proteome also show conservation across taxa (Bayram et al. 2016; Pini et al. 2025). For instance, Sperm-leucylaminopeptidases (S-Laps), first identified in the *Drosophila melanogaster* sperm proteome, have been identified in many insects (Dorus et al. 2011; Whittington et al. 2015; Whittington et al. 2017; Bayram et al. 2019; Degner et al. 2019; Laurinyecz et al. 2019; Kumar et al. 2022; Gomez et al. 2024). Similarly, peptidases and peptidase inhibitors are common features of seminal fluid proteomes that initiate cascades of downstream interactions between female and male proteins to ensure long-term postmating responses (LaFlamme et al. 2014; Avila et al. 2015; Singh et al. 2018; Plakke et al. 2019; Wainwright et al. 2021). Studying the seminal proteome in non-model organisms allows testing commonly held assumptions about reproductive proteins (Zuk et al. 2014).

The common bedbug (*Cimex lectularius*) presents an attractive model to study the factors shaping the seminal proteome, with unique selective pressures potentially acting on the male ejaculate. First, in species where males are the heterogametic sex (i.e., XY), genes with male-biased expression are expected to be underrepresented on the X chromosome which spends two-thirds of time in females (Vibranovski et al. 2009). The karyotype in bedbugs is 2n = 26 + X_1_X_2_Y in males (Slack 1939; Ueshima 1967; Grozeva et al. 2010), which may reduce sex biased selection and alter the landscape of sex biased genes. Second, bedbugs obligately mate by traumatic insemination, where the male intromittent organ (paramere) pierces the female cuticle, bypassing the female reproductive tract, which is used only during oviposition (Siva-Jothy 2006). Insemination takes place in the mesospermalege, a novel organ that has evolved in the Cimicidae (Roth et al. 2024), which reduces mating costs to females (Morrow and Arnqvist 2003; Reinhardt et al. 2003). As the site of insemination, sperm first interact with the female in the mesospermalege before sperm exit into the female haemolymph (Usinger 1966; Martens et al. 2026). Sperm then travel through the haemolymph and cross into the female reproductive tract to enter the oviduct via an as yet unknown mechanism, where some sperm are stored in a unique storage organ (Usinger 1966). Other sperm move up the oviduct and, unlike in other insects, fertilise eggs in the ovaries (Usinger 1966). Sperm must therefore traverse multiple tissues, membranes and substrates to reach the egg (Usinger 1966).

To explore features of the bedbug seminal proteome associated with traumatic insemination, we provide a detailed proteomic investigation of the male bedbug reproductive organs, including the sperm and seminal proteomes. This is the first high-throughput seminal proteome of a hemimetabolous insect, greatly increasing the number of ejaculate proteins in bedbugs (Reinhardt, Wong, et al. 2009). Recently published genomic resources (Law et al. 2025; Miles et al. 2025) allow us to further investigate the evolutionary dynamics of bedbug reproductive proteins, including rates of molecular evolution and chromosomal distribution. Finally, we compare the bedbug seminal proteome to *Drosophila* and mosquitoes (another blood-feeding insect). In doing so, we identified an expansion of the *S-Lap* gene family in the Cimicomorpha, showing retention of catalytic activity based on sequence similarity and *in silico* protein-ligand interactions.

## RESULTS

Label-free quantitation using LC-MS/MS of the *Cimex lectularius* male reproductive tract tissues and sperm provide a unique and quantitative view of the bedbug seminal proteome. In total, we identified 4,651 proteins, of which we retained 3,348 (71.7%; Table 1; Fig. S2) after filtering proteins identified by fewer than 2 unique peptides and default filtering steps in *MSstats* (Kohler et al. 2023).

**Table 1.** Total numbers of proteins identified in each tissue/sperm and the number of tissue/sperm-biased proteins (as a percentage of total in brackets. SFCs, seminal fluid containers.

|  | Testes | Sperm | Mesadenes | SFCs |
| --- | --- | --- | --- | --- |
| No. proteins identified | 3244 | 2494 | 3017 | 2272 |
| No. tissue-biased (%) | 535 (16.5%) | 487 (19.5%) | 708 (23.5%) | 239 (10.5%) |

Unlike most other insects, true bugs have separate organs which are the source of seminal fluid production; the mesadenes, which are physically separated from the seminal fluid storage organ, the seminal fluid containers (Fig. 1A). A principal component analysis (PCA) of normalised protein abundances showed the first two PCs explained more than 83% of variation, separating all four tissues (Fig. 1B). PC1 (47.9% variance explained) separated the sperm and testes from the seminal fluid containers and mesadenes (Fig. 1B). PC2 (35.2% variance explained) further separated the sperm and seminal fluid containers from the testes and mesadenes. This suggests some gross similarities between the organs that are the sources of protein production (testes and mesadenes), separate from the sinks (sperm and seminal fluid containers).

**Figure 1.**
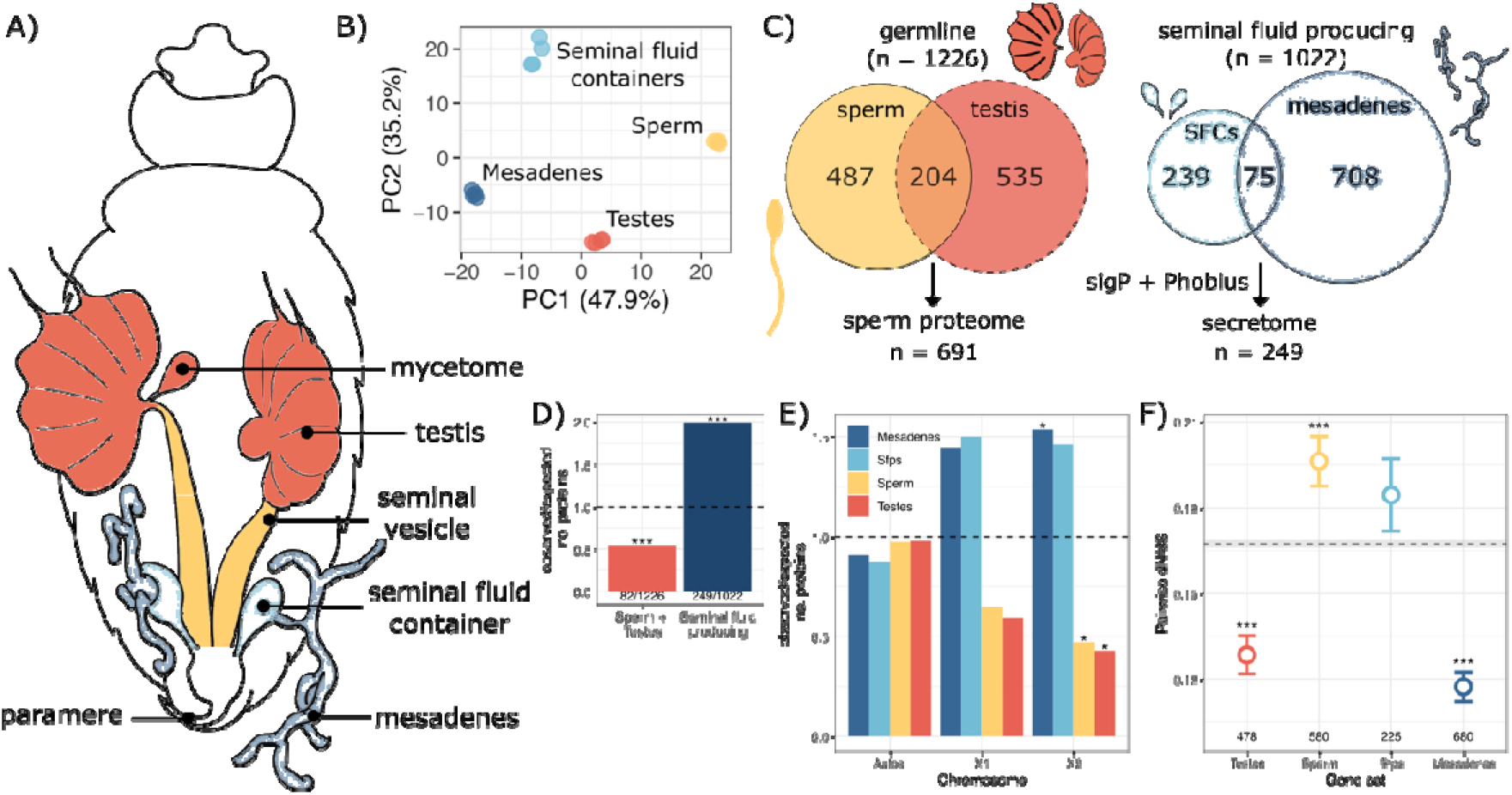
**(A)** Dorsal view of male Cimex lectularius reproductive system highlighting organs dissected for proteomics (adapted from (Usinger 1966)). Sperm were isolated from the seminal vesicles. Bacteriomes (aka mycetomes) were included with the testes for LC-MS/MS. **B)** Principal component analysis of protein abundances for the 500 most variable proteins. **C)** Reproductive protein classification workflow. Numbers in Venn diagrams show number of proteins biased towards each tissue (log2FC > 1 & adjusted p-value < 0.05). **D)** Enrichment of proteins containing a signal peptide sequence in the bedbug reproductive proteome. Numbers below bars are the observed and expected number of proteins in each group, respectively. The dashed line indicates the null expectation, i.e., the expected proportion of proteins in the whole proteome. P-values obtained from Χ^2^ tests. **E)** Chromosomal distribution of male-biased proteins on the autosomes and each of the two C. lectularius X chromosomes. Asterisks represent results from comparing the observed to expected number of proteins on each chromosome after multiple testing correction using X^2^ tests. The dashed line indicates the null expectation. **F)** Pairwise nonsynonymous (dN) to synonymous (dS) nucleotide substitution rate, dN/dS (mean ± standard error) between C. lectularius and C. hemipterus. Numbers below points indicate numbers of genes in each category. Asterisks represent results from Mann-Whitney U tests comparing each gene set to the genome average shown by dashed line (dN/dS = 0.167 ± 0.001, n = 17580). * p < 0.05, ** p < 0.01, *** p < 0.001. Sfp, seminal fluid proteins. SFCs, seminal fluid containers.

### Characterisation of the male reproductive proteome

We defined proteomes for each tissue/sperm based on proteins having higher abundance (or the exclusive presence) compared to all other tissues (log2FC > 1 & adjusted *p*-value < 0.05; Fig. 1C; Table 1; Fig. S2&S3).

#### Testis and sperm proteome

The combined testis and sperm proteome comprised 1,226 proteins and showed underrepresentation of proteins containing a signal peptide sequence (Χ^2^ = 30.67, df = 1, *p* < 0.001; Fig. 1D). Sperm- and testes-biased proteins were underrepresented on chromosome X_2_ (sperm: Χ^2^ = 10.0, df = 1, *p* = 0.046; testes: Χ^2^ = 9.14, df = 1, *p* = 0.05; Fig. 1E & Fig. S5).

The putative sperm proteome, comprising proteins biased towards sperm or sperm and testes (n = 691; Fig. 1C) was evolving at an elevated rate compared to the genome average (mean pairwise dN/dS ± standard error = 0.20 ± 0.01, n = 580; pairwise Wilcoxon test, *p* < 0.001; Fig. 1F). The sperm proteome showed GO enrichment of biological processes involving metabolic and glycolytic processes, proteolysis, cilium assembly and microtubule-based process; cellular components typically associated with sperm cells, such as mitochondria, cilium, axoneme, dynein complex, cytoplasm and microtubules; and molecular functions including ion binding and serine-type endopeptidase activity (Table S3).

Proteins exclusively biased towards testes (n = 535) showed GO enrichment of biological processes including protein folding, cellular transport and organisation and translation; cellular components including proteasomes, cytoplasm, microtubules and mitochondria; molecular functions including protein folding, ATP binding and heat shock protein binding (Table S4). Endosymbiotic bacteria (e.g., *Wolbachia*) (Thongprem et al. 2020) are localised in bacteriomes (aka mycetomes) attached to the *vas deferens* at the base of the testes (Usinger 1966; Hosokawa et al. 2010). As expected, *Wolbachia* proteins showed the highest abundance in the testes/bacteriome samples (Fig. S4).

#### Mesadene and seminal fluid proteome

The seminal fluid producing proteome, including proteins biased towards mesadenes and/or seminal fluid containers comprised 1022 proteins (Fig. 1C). Proteins containing a signal peptide sequence were overrepresented in the mesadenes and seminal fluid containers (Χ^2^ = 123.45, df = 1, *p* < 0.001; Fig. 1D). We classified the 249 proteins containing a signal peptide sequence from the seminal fluid producing organs as our putative seminal fluid proteome (Fig. 1C); a number similar to the number of “high confidence” Sfps identified in *Drosophila melanogaster* (Wigby et al. 2020). While the presence of a signal peptide sequence is indicative of a protein being secreted (Teufel et al. 2022) and a hallmark of seminal fluid proteins, not all secreted proteins contain signal peptides. We found a subset of putative seminal fluid proteins expressed highly (in the top 20% most abundant proteins) in sperm (3.6%; 9/249) and testes (8%; 20/249), in line with recent studies suggesting a more complex landscape of seminal fluid biology (Garlovsky et al. 2022; McCullough et al. 2022).

Seminal fluid proteins were evolving at a similar rate to sperm (*p* = 0.106) but were not evolving faster than the genome average (dN/dS = 0.18 ± 0.01, n = 224; *p* = 0.135; Fig. 1F). The seminal fluid proteome showed GO enrichment for biological processes including protein folding and proteolysis, glycosylation and negative regulation of peptidase activity; cellular components including endoplasmic reticulum and extracellular space; and molecular functions including ion and molecule binding, and (serin-type) endopeptidase inhibitor activity (Table S5).

The remaining 773 proteins classified as the seminal fluid producing mesadenal category were overrepresented on X_2_ (Χ^2^ = 11.8, df = 1, *p* = 0.035; Fig. 1E). Mesdadenal proteins showed GO enrichments often associated with accessory gland/secretory functions; biological processes such as translation, protein transport and Golgi/vesicle transport; cellular components including ribosome, endoplasmic reticulum and Golgi apparatus; molecular functions including RNA- and GTP-binding (Table S6).

### Comparison with other seminal proteomes

In the bedbug sperm proteome (n = 691), 189 orthologs of *D. melanogaster* sperm proteins were identified (Garlovsky et al. 2022; McCullough et al. 2022) and 58 *Aedes aegypti* sperm proteins (Degner et al. 2019). Among the most abundant proteins in the bedbug sperm proteome were orthologs of *D. melanogaster* sperm proteins, including proteins involved in spermatid development (e.g., betaTub56D, bb8), S-Laps, mitochondrial proteins (e.g., P5CDh1, Cs1, Ant2, mAcon, Mitofilin) and Y-linked fertility factors (e.g., ORY, WDY, Ppr-Y, kl-3, kl-5) (Table 2; Table S7). The bedbug seminal fluid proteome (n = 249) contained fewer orthologs; just 15 *D. melanogaster* seminal fluid proteins (Wigby et al. 2020), and 5 *Ae. aegypti* seminal fluid proteins (Degner et al. 2019).

**Table 2.**
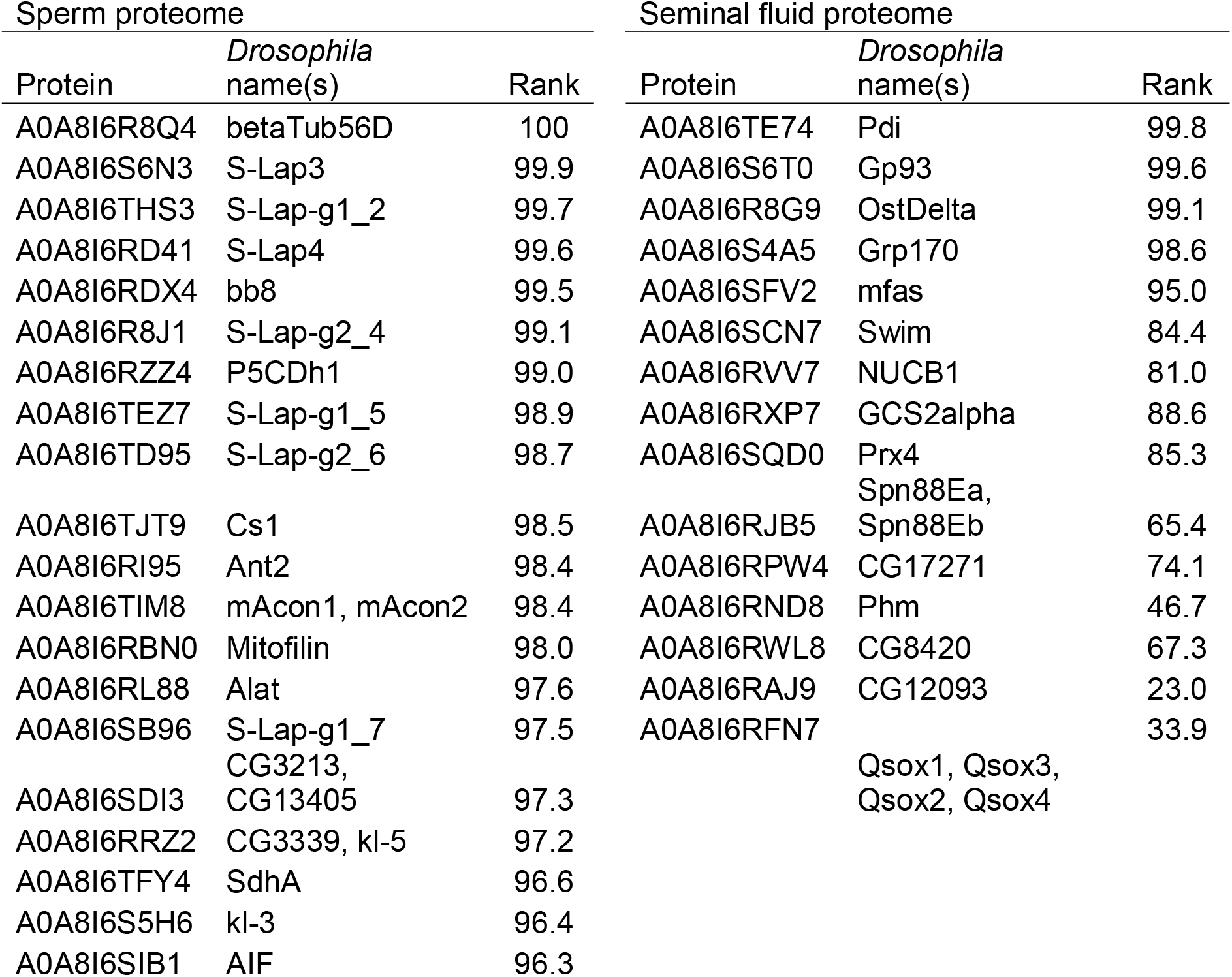
The most abundant proteins in the bedbug sperm- and seminal fluid-proteomes with D. melanogaster orthologs (rank ordered by abundance). Bedbug proteins matching more than one Drosophila gene are separated by a comma. S-Laps are labelled based on one-to-one orthology with D. melanogaster or membership to S-Lap cluster 1 (g1) or 2 (g2) (Dorus et al. 2011) followed by a unique ID. D. melanogaster Sfps are noted with an asterisk (all found in the Sfp category).

### Sperm-leucylaminopeptidase evolution

Notably, 6 of the 10 most abundant bedbug sperm proteins with *D. melanogaster* orthologs were S-Laps (Fig. 2; Table 2) (Dorus et al. 2011). We identified 13 *C. lectularius* S-Lap orthologs in total; five belonging to S-Lap cluster 1, eight in cluster 2 (Dorus et al. 2011), all of which were most abundant in sperm (Fig. S6). To further explore orthology between the bedbug and *Drosophila* S-Laps we used BLASTp (Camacho et al. 2009) which showed matches to all 8 *Drosophila* S-Laps. A more stringent reciprocal best BLAST search (e-value = 1e-2, min. ident = 30, min. coverage = 50) returned four one-to-one S-Lap orthologs: S-Laps 2, 3, 4 and 7 (Fig. 3B).

**Figure 2.**
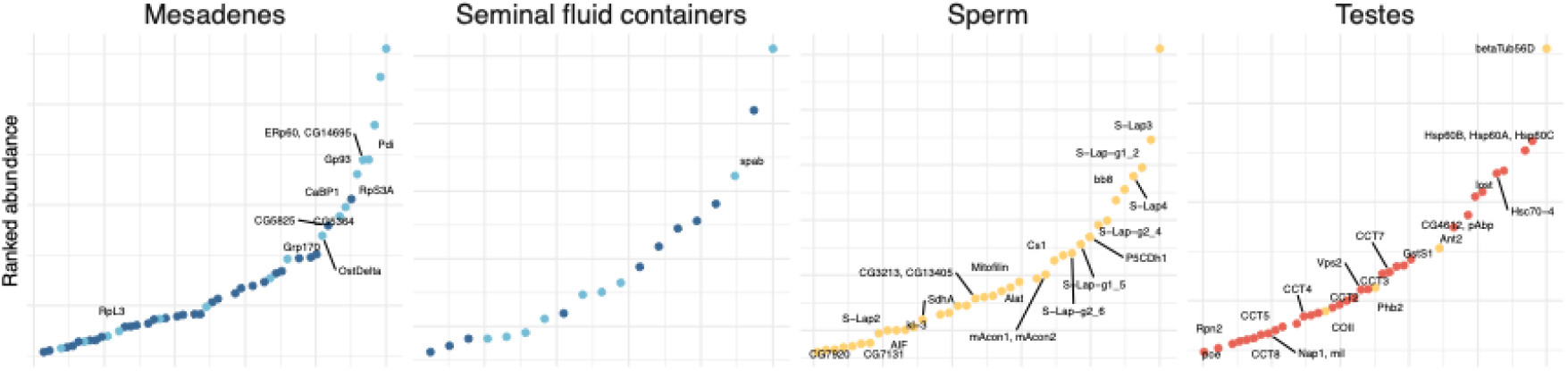
Ranked abundance of the top 5% most abundant proteins in each tissue/sperm. Points are coloured by proteome category (mesadenes, dark blue; seminal fluid proteins, light blue; sperm, yellow; testes, orange). Orthologs of Drosophila melanogaster sperm (Garlovsky et al. 2022) and seminal fluid proteins (Wigby et al. 2020) are labelled. S-Laps are labelled based on one-to-one orthology with D. melanogaster or membership to S-Lap cluster 1 or 2 (Dorus et al. 2011) followed by a unique ID.

**Figure 3.**
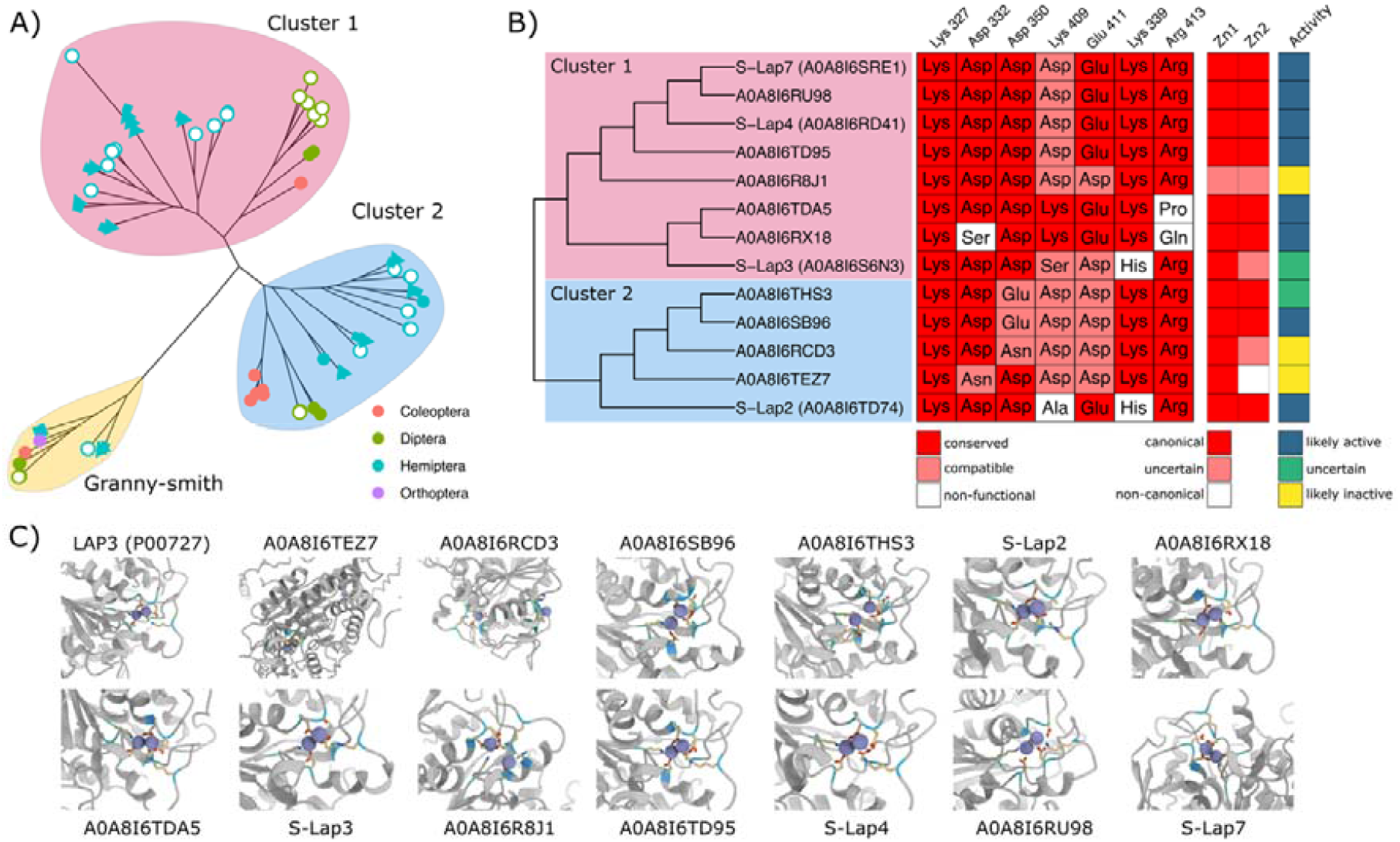
Sperm-leucylaminopeptidase (S-Lap) enzymatic activity in bedbugs. **A)** Unrooted gene tree of S-Lap and granny-smith orthologs in insects. Protein sequences were aligned using MAFFT (Madeira et al. 2024) and bootstrapped 1,000 times. Tip colour indicates Order and shape indicates species. Drosophila and bedbug S-Laps shown with open circles. **B)** Alignment of bedbug S-Lap orthologs focused on amino acid residues against the M17 amino peptidase metal binding motif indicated above (Burley et al. 1992). Amino acids are coloured by alignment to consensus (conserved, red; compatible, pink; non-functional, white). Zn^2+^ ion binding at the “loose” binding site (Zn1) and “tight” binding site (Zn2) is scored as canonical (red), uncertain (pink) or non-canonical (white). Overall activity is scored as likely active (blue), uncertain (green), or likely inactive (yellow) based on coordinating residue geometry, number and inter-ligand angles and inter-zinc distances measured from AlphaFold3 models in ChimeraX. **C)** AlphaFold predictions for structures of the canonical M17 LAP, cytosol aminopeptidase (LAP3; UniProt ID: P00727), and bedbug S-Laps focussed on zinc binding sites. Residues containing atoms within 4Å of Zn^2+^ ions (purple spheres) are highlighted: N, blue; O, red; S, yellow; C, blue.

There appear to be no S-Lap orthologs in more distantly related Hemiptera (*Acyrthosiphon pisum*) or Orthoptera (*Gryllus bimaculatus*), but S-Lap orthologs were present in the Coleoptera (*Callosobruchus maculatus*) (see also (Gomez et al. 2024) (Fig. 3A). However, orthologs of *granny-smith*, the closest S-Lap paralog (Dorus et al. 2011), were identified in all species studied (Fig. 3A & Fig. S7). Across the 73 S-Lap orthologs from 7 species studied (including *D. melanogaster*), proteins first cluster by S-Lap cluster (1 or 2) rather than species based on sequence similarity, suggesting the origin of S-Laps predates the separation of the Holometabola from the Hemimetabola (Misof et al. 2014).

S-Laps belong to the M17 family of leucyl aminopeptidases (Dorus et al. 2011) which bind two Zn^2+^ ions (Maric et al. 2009). However, in *D. melanogaster*, S-Laps belonging to “cluster 1” have mutations at most of the active site residues and sperm display very weak enzymatic activity (Dorus et al. 2011; Laurinyecz et al. 2019). Most (11 out of 13) bedbug S-Laps, including those in “cluster 1”, retain at least 4 out of 5 metal ion binding residues including all the residues coordinating tight Zn^2+^ binding and cofactor binding sites (Fig. 3B).

Structure-based predictions to determine 3D coordination of Zn^2+^ ions also showed that most bedbug S-Laps (at least 8 out of 13) retain the M17 LAP zinc binding coordination geometry (Fig. 3B,C). However, five bedbug S-Laps (two in cluster 1, three in cluster 2) are likely inactive, with inter-zinc distances exceeding the ∼3Å observed in the conserved M17 structure (Burley et al. 1992; McGowan et al. 2010) (Fig. 3B,C).

This pattern of (i) retention of metal binding active site residues, and (ii) 3D coordination of Zn^2+^ was observed across the species we studied outside of *Drosophila* including other hemipterans and hemimetabolous insects as well as in Diptera and Coleoptera (Fig. S7). Notably, *C. maculatus* S-Laps in “cluster 1” do not have the conserved Asp residue coordinating site 2 tight zinc binding and lack zinc binding geometry (Fig. S7).

## DISCUSSION

Bedbugs are unusual among insects in obligately mating by traumatic insemination. After insemination, sperm move in aggregations out of the mesospermalege towards the ovaries or female sperm storage organs (Usinger 1966; Martens et al. 2026). Sperm must therefore cross multiple tissue membranes, likely shaping key adaptations in the bedbug seminal proteome. Our analysis revealed conserved and unique features of the sperm and seminal fluid proteome with possible links to this unusual sperm behaviour.

Sperm and testes proteins were underrepresented on chromosome X_2_, as expected for proteins with male-biased expression (Vibranovski et al. 2009). However, the same pattern was not found for chromosome X_1_. The two X chromosomes in *Cimex* are thought to be derived from fission of an ancestral X (Grozeva et al. 2010; Poggio et al. 2014). Mesadenal proteins, on the other hand, were overrepresented on X_2_, which may result if new mesadenal genes have arisen on the X (Levine et al. 2006). The multiple sex chromosomes present interesting opportunities for unusual evolutionary dynamics that warrant further investigation.

We did not observe the commonly found pattern of rapidly evolving seminal fluid proteins (Swanson and Vacquier 2002), but found few seminal fluid protein orthologs in *Drosophila* or *Aedes* mosquitoes. Surprisingly, elevated rates of molecular evolution were observed in the bedbug sperm proteome. In many taxa, the sperm proteome evolves slowly, thought to result from purifying selection (Garlovsky et al. 2022). Elevated rates of sperm evolution might be driven by sexually antagonistic co-evolution in the bedbug associated with traumatic insemination and movement of sperm through the haemolymph. The mating rate in bedbugs is at least ten times above the female optimum (Stutt and Siva-Jothy 2001; Reinhardt et al. 2011), setting the stage for sexual conflict affecting ejaculate × female interactions (Morrow and Arnqvist 2003; Reinhardt et al. 2003; Sirot et al. 2015). Sperm traversing the female haemolymph and female reproductive tract might entail novel and unusual selective pressures acting on sperm and seminal fluid proteins. As such, it is unclear if or for how long seminal fluid associates with sperm after insemination, as found in other species (Misra and Wolfner 2020). We also found a subset of putative seminal fluid proteins were abundant in sperm and testes, in line with recent studies which point to a possible testis origin of some seminal fluid proteins (Garlovsky et al. 2022; McCullough et al. 2022).

Despite overall elevated rate of evolution in the bedbug sperm proteome, many of the most abundant sperm proteins had orthologs in other insects. Most notably, we identified 13 Sperm-leucylaminopeptidases (S-Laps) in bedbugs and other insects including other hemimetabolous insects. The S-Laps in *Drosophila* show evidence of purifying selection (Dorus et al. 2011) and indeed were evolving more slowly than the rest of the genome in bedbugs (Fig. S8). Other S-Lap orthologs have been identified in Lepidoptera, Coleoptera and Diptera, suggesting an ancient origin of these sperm proteins after more than 380 million years since the last common ancestor (Dorus et al. 2011; Whittington et al. 2015; Whittington et al. 2017; Bayram et al. 2019; Degner et al. 2019; Laurinyecz et al. 2019; Garlovsky et al. 2022; Kumar et al. 2022; Gomez et al. 2024).

S-Laps belong to the M17 family of metal-dependent leucyl aminopeptidases, but in *Drosophila* have mostly lost enzymatic activity (Dorus et al. 2011; Laurinyecz et al. 2019). Our functional analysis revealed that S-Laps in *Cimex* retain the Zn^2+^ binding motif, suggesting that S-Laps retained enzymatic properties. Additional direct biochemical assay of *Cimex* S-Laps will help support the idea that enzymatic activity has been secondarily lost in *Drosophila*. Our analysis suggests neofunctionalisation of this class of M17 leucyl-aminopeptidases while retaining an active site in bedbugs and other insects. S-Laps have been identified in whirligig beetles and also retain the metal binding motif (Gomez et al. 2024). Whirligig sperm move in conjugates attached to spermatostyles (Breland and Simmons 1970). Bedbug sperm also move in conjugates, lending support to the notion that S-Laps play a role in sperm conjugation and cooperation (Gomez et al. 2024).

### Conclusions

In conclusion, our study provides a substantially expanded database of male reproductive proteins in bedbugs using high-throughput proteomics, increasing coverage of the bedbug seminal proteome by two orders of magnitude (Reinhardt, Wong, et al. 2009). Our analysis provides insights into the landscape of sperm and seminal fluid dynamics in an insect with unique features of reproductive anatomy and physiology. The peculiarities of traumatic insemination and multiple sex chromosomes in the bedbug have led to unique evolutionary characteristics of the seminal proteome. Sperm genes are underrepresented on the second X chromosome and evolve rapidly. S-Laps have retained catalytic activity, which may aid in sperm conjugation. A better understanding of the proteins important for reproduction will aid the development of effective strategies to control and suppress populations of insect pests and disease vectors (Doggett et al. 2012). Unlike many other blood-feeding insects, bedbugs do not transmit any known human pathogens (Doggett et al. 2012). Nevertheless, bedbugs are an expanding global pest, causing sleeplessness, skin irritation, severe allergic reactions and growing economic costs (Reinhardt and Siva-Jothy 2007; Reinhardt, Kempke, et al. 2009; Doggett et al. 2012). Our results provide a useful resource for future studies and a list of candidate proteins involved in reproduction in this economically important insect.

## MATERIALS AND METHODS

### Bedbug maintenance

We used bedbugs originally collected in London, UK (F4 population) in 2006 and maintained at TU Dresden, Germany, as a large, outbred population with overlapping generations. Bedbugs were maintained at 26ºC, 70% relative humidity on a 12:12 hour light:dark cycle. Bedbugs were fed on human blood once each week during the juvenile stages following long-term experimental protocols (Reinhardt et al. 2003). We used males aged 3 weeks post-final moult. Males were housed in groups of 10-15 and fed once a week for 2 weeks after eclosion as adults and left to mature for a further week.

### Tissue collection and protein extraction

All dissections were performed in freshly prepared ice-cold phosphate-buffered saline (1X PBS) with 1X protease inhibitors (HALT) and 5mM EDTA (Thermo Fisher). We anaesthetised males on ice and removed reproductive tracts with forceps, insect pins, and dissecting scissors under a stereo dissecting microscope. We had four replicates consisting of 4-6 males each.

To collect sperm samples, seminal vesicles were separated from the remaining reproductive tract tissue and transferred to a fresh drop of PBS/HALT. We then made a small incision at the distal end of the seminal vesicle and exuded the sperm mass using insect pins. The sperm mass was then transferred to an Eppendorf containing 1mL PBS/HALT kept on ice. Seminal fluid containers and mesadenes were moved to a fresh drop of PBS/HALT and separated using insect pins. Mesadenes were transferred to an Eppendorf containing 1mL PBS/HALT kept on ice. Seminal fluid containers were transferred in 2 µL PBS/HALT per gland to a separate Eppendorf. Testes were separated from the remaining tissue using insect pins and transferred to an Eppendorf containing 1mL PBS/HALT kept on ice. Sperm, testis, and mesadene samples were pelleted at 15,000 × G for 2 minutes at 4ºC, supernatant carefully removed, and the pellet resuspended by addition of 1 mL PBS/HALT followed by an additional 2 mins centrifugation at 15,000 × G. The washing procedure was repeated twice, and the final pellet resuspended in 1 mL PBS/HALT and stored at -80ºC. Seminal fluid containers were freeze/thawed and vortexed for 30 seconds three times and then spun at 15,000 × G for 5 minutes at 4ºC and stored at -80ºC suspended in 10 µL of PBS/HALT.

### Liquid chromatography tandem mass spectrometry

All LC-MS analyses were performed at the Biosciences Mass Spectrometry Core Facility (https://cores.research.asu.edu/mass-spec/) at Arizona State University. For LC–MS/MS, solubilised proteins were quantified (Thermo Fisher EZQ Protein Quantitation Kit or the Pierce BCA). Proteins were reduced with 50□mM dithiothreitol (Sigma-Aldrich) at 95□°C for 10□min and alkylated for 30□min with 40□mM iodoacetamide (Pierce). Proteins were digested using 2.0□μg of MS-grade porcine trypsin (Pierce), and peptides were recovered using S-trap Micro Columns (Protifi) per manufacturer directions. Recovered peptides were dried via speed vac and resuspended in 30 µL of 0.1% formic acid. All data-dependent mass spectra were collected in positive mode using an Orbitrap Fusion Lumos mass spectrometer (Thermo Scientific) coupled with a Dionex UltiMate 3000 system equipped with an NCS-3500 nano capillary pump and UltiMate 3000 RS autosampler. One µL of the peptide was fractionated using an Easy-Spray LC column (50□cm□Å ∼75□μm ID, PepMap C18, 2□μm particles, 100□Å pore size, Thermo Scientific). Electrospray potential was set to 1.9□kV and the ion transfer tube temperature to 275□°C. The mass spectra were collected using the method optimised for peptide analysis provided by Thermo Scientific. Full MS scans (350–1500□m/z range) were acquired in profile mode with the following settings: Orbitrap resolution 120,000, cycle time 3□s and mass range “Normal;” RF lens at 35%, and the AGC set to “Standard”. Monoisotopic peak determination (MIPS) at “peptide” and included charge states 2– 7; dynamic exclusion at 60□s, mass tolerance 10 ppm, intensity threshold at 5.0e4; MS/MS spectra acquired in a centroid mode using quadrupole isolation window at 1.6 (m/z); Higher-energy collisional dissociation (HCD) energies at 35%. Spectra were acquired over a 180-min gradient, flow rate 0.250 µL/min as follows: 0–10□min at 0-2%, 10–100□min at 2–8%, 100–110□min at 8–32%, 110-120□min at 32–35%, 120–125□min at 35–45%, 125–155 min at 45-98%,155–165 min at 98%, 165 – 180□min at 2%,followed by post run wash step.

### Protein identification and label-free quantification

We analysed raw files searched against the UniProt (www.uniprot.org) *Cimex lectularius* database that included two bacterial predicted proteomes (CIMLE_UP000494040; wCle UP000031663; Rickettsia UP000826664) using Proteome Discover 2.5 (Thermo Scientific). Input data included Trypsin as enzyme, maximum missed cleavage site 3, min/max peptide length 6/144, and precursor ion (MS1) mass tolerance set to 20 ppm and fragment mass tolerance set to 0.5 Da. Carbamidomethyl (C) was specified as fixed modification, and dynamic modifications set to Aceytl and Met-loss at the N-terminus, and oxidation of Met. SequestHT and Percolator ID validation was performed using a minimum of 1 peptide identified using a concatenated target/decoy strategy and false-discovery rate (FDR) set to 1.0%. The Minora feature detector identified between run peak matches and quantification intensities, normalised and scaled to a maximum of 100. Intensity values were averaged across all six runs and ranked from highest to lowest to obtain a relative abundance profile. A combined list of predicted signal peptide sequences from the entire *C. lectularius* proteome was obtained using signal-P 6 (Teufel et al. 2022) and Phobius (Käll et al. 2004).

### Ortholog identification

We downloaded proteomes for *Cimex hemipterus*, assassin bug (*Triatoma infestans*), kissing bug (*Rhodnius prolixus*), pea aphid (*Acyrthosiphon pisum*), seed beetle (*Callosobruchus maculatus*), yellow fever mosquito (*Aedes aegypti*) and fruit fly (*Drosophila melanogaster*) from uniprot.org and two-spotted cricket (*Gryllus bimaculatus*) from NCBI (https://www.ncbi.nlm.nih.gov/) (Fig. S1; Table S1). Species were chosen because of their evolutionary relationship to *Cimex lectularius*, i.e., the true bugs and hemimetabolous insects (aphid, cricket), because of similar biology, such as blood feeding (mosquito), or because of a wealth of data on reproductive biology, including the occurrence of copulatory wounding (*D. melanogaster* and seed beetle). Hierarchical orthologs were identified in each species with *OrthoFinder* using default settings (Emms and Kelly 2015; Emms and Kelly 2019).

### Evolutionary dynamics

We obtained previously calculated pairwise rates of nonsynonymous (dN) and synonymous (dS) nucleotide substitution rates (dN/dS) and chromosomal positions of genes (Martens et al. 2026). Briefly, rates of molecular evolution (dN/dS) were calculated between *C. lectularius* and *C. hemipterius* using PAML (Yang 2007). Chromosomal distribution of genes was determined by aligning coding sequences from the *C. lectularius* reference genome Clec_2.1 (Benoit et al. 2016) to the recently published chromosome-level *C. lectularius* genome assembly (Miles et al. 2025) using BLASTn (Camacho et al. 2009).

### Gene ontology enrichment analysis

We performed gene ontology enrichment analysis using the topGO package (Rahnenfuhrer 2022) in R (R Core Team 2024) using the GO annotations from uniprot.org. The entire bedbug proteome was used as background for enrichment tests. Statistical analyses in topGO used Fisher’s exact tests with topnodes = 10, and corrected p-values for multiple testing using the FDR method (Benjamini and Hochberg 1995). All GO terms reported had an FDR-corrected *p*-value < 0.05.

### Statistical analysis

All statistical analysis was performed in R v4.5.0 (R Core Team 2024). Output from Proteome Discoverer was imported into R and formatted for use in the *MSstats* pipeline (Kohler et al. 2023) using the following options: normalisation = quantile, imputed missing values using the TMP method, MBimpute = TRUE.

To identify proteins biased towards each tissue (seminal fluid containers, mesadenes, testes) or sperm, we performed pairwise differential abundance analysis between each tissue/sperm. We considered a protein to be biased towards a tissue/sperm based on a log2-fold-change > 1 and false discovery corrected *p*-value < 0.05 compared to all other tissues. Proteins had to be identified in two or more replicates of a tissue/sperm to count for that tissue/sperm. We then classified proteins into 1 of 4 categories: (1) the putative seminal fluid proteome was defined as proteins biased towards the mesadenes and/or seminal fluid containers containing a signal peptide sequence; (2) the remaining proteins from the seminal fluid producing organs were classified as the mesadenal (seminal fluid producing) proteome; (3) the putative *C. lectularius* sperm proteome was defined as proteins more abundant in sperm or sperm and testes; (4) the remaining proteins biased towards testes we classified as the testes proteome (Fig. 1C).

### Predicting protein-interactions

We characterised catalytic activity for S-Laps based on conservation of the M17 aminopeptidase metal-binding and catalytic residue motif (Maric et al. 2009). Orthologs of S-Laps and granny-smith (UniProt ID: Q9V3D8) identified using *OrthoFinder* and cytosol aminopeptidase (LAP3; UniProt ID: P00727) were aligned using *MUSCLE* (Edgar 2022). The catalytic domain of each sequence was identified by mapping a ±150 residue window around the active site centre of *granny-smith*. The position equivalent to the start of the motif (Lys327) was pinned in each sequence and the 7 active site positions scored at fixed offsets from the anchor. Each position was classified as (i) conserved, exact match to the M17 LAP consensus; (ii) compatible, a residue with structural or mutagenesis evidence for metal coordination in M17 LAPs (Burley et al. 1992; Gu and Walling 2002; Dorus et al. 2011); or (iii) non-functional (Table S2).

Zinc coordination sites were also assessed using mmCIF results from modelling complexes of each protein (after removing signal peptide sequences identified in *Phobius*) with two Zn^2+^ ions using Alphafold3 (Abramson et al. 2024). Zinc coordination geometry was assessed with ChimeraX v.1.11.1 (Meng et al. 2023) in headless mode using a custom Python script. Atoms within 3.5Å of each zinc ion were identified and contacts classified as canonical if they involved the sidechain nitrogen of lysine (NZ) or carboxylate oxygens of aspartate (OD1/OD2) or glutamate (OE1/OE2) consistent with the experimentally derived coordination chemistry of M17 leucyl aminopeptidases (PDB: 1LAP) (Burley et al. 1992). Where both carboxylate oxygens of a single Asp or Glu residue fell within the distance cutoff (bidentate geometry) only the closer oxygen was retained.

Each zinc ion was classified as occupying either the exchangeable site (Zn1, characterised by lysine coordination) or the tight site (Zn2, characterised by two aspartate contacts and absence of lysine), based on coordination chemistry of LAP3 (P00727). We applied a scoring rubric (from 1 to 10) for coordination environments of Zn1 and Zn2 that incorporated presence of canonical coordinating residues (6 points), coordination number (2 points), bond distance (1 point, penalised for contacts > 3.2Å) and mean pLDDT confidence at the binding site (≥ 70, 1 point). Each binding site was classified as canonical (score ≥ 6), uncertain (score 3-5), or non-canonical (score < 3). Distance thresholds were set to account for the systematic overestimation of metal-ligand bond lengths in AF3 predictions relative to experimental crystal structures.

After scoring each site, structures were assessed for presence of a shared bridging ligand/residue providing sidechain contacts to both zinc ions and Euclidean distance between the two zinc ions was calculated to determine if zinc ions resided in the same zinc pocket (≤4Å) (Burley et al. 1992). Proteins were classified as; “likely active”: both binding sites canonical and zinc ions residing within the same zinc pocket; “uncertain”: both canonical sites and both zincs within the inter-zinc distance but no bridging ligand; or “likely inactive”, i.e., non-functional: one or two non-canonical binding sites or separate binding pockets (inter-zinc distances > 7Å).

## ACKNOWLEDGEMENTS

We would like to thank Christin Froschauer and Susanne Broschk for help in the lab and Caitlin McDonough-Goldstein for comments on an earlier version of the manuscript. We thank members of the ASU Biosciences Core Mass Spectrometry facility, Kyle Tucker, Sydney Canning and Yishai Gilron for the acquisition of mass spectra and downstream computational analyses. Yan Ge and the ZIH team at TU Dresden provided valuable assistance in implementing AlphaFold3. This work was funded by DFG grants to OO 521/4-1 and KR 1666/4-1. We are grateful for high-performance computing at the NHR Center of TU Dresden, jointly supported by the Federal Ministry of Education and Research and the state governments participating in the NHR (www.nhr-verein.de/unsere-partner).

