## Supplementary material for "Evolution and adaptations of the seminal proteome in an insect with traumatic insemination"

Martin D. Garlovsky^1*^, Oliver Otti^1^, Klaus Reinhardt^1^, Timothy L. Karr^2,3^

**AFFILIATIONS**

^1^Applied Zoology, Faculty Biology, Technische Universität Dresden, Dresden, Germany

^2^Biosciences Mass Spectrometry Core Research Facility, Knowledge Enterprise, Arizona State University, USA

^3^Neurodegenerative Disease Research Center, The Biodesign Institute, Arizona State University, USA

**ACKNOWLEDGEMENTS**

We would like to thank Christin Froschauer and Susanne Broschk for help in the lab and Caitlin McDonough-Goldstein for comments on an earlier version of the manuscript. We thank members of ASU Biosciences Core Mass Spectrometry facility Kyle Tucker, Sydney Canning and Yishai Gilron for acquisition of mass spectra and downstream computational analyses. Yan Ge and the ZIH team at TU Dresden provided valuable assistance implementing AlphaFold3. This work was funded by DFG grants to OO 521/4-1 and KR 1666/4-1. We are grateful for high-performance computing at the NHR Center of TU Dresden, jointly supported by the Federal Ministry of Education and Research and the state governments participating in the NHR ([www.nhr-verein.de/unsere-partner](http://www.nhr-verein.de/unsere-partner)).

**DATA AVAILABILITY**

Data and details of the analysis will be made available on the Open Access Repository of Saxon Universities (<https://opara.zih.tu-dresden.de/home>) and GitHub (https://github.com/MartinGarlovsky/Bedbug_proteomics). Proteomic data have been deposited to the ProteomeXchange Consortium via the PRIDE partner repository (Perez-Riverol et al. 2022) with the identifier PXD075584.

**SUPPLEMENTARY METHODS**

***Table S1.*** Number of proteins in each species

| **Species** | **No. proteins** | **No. proteins in orthogroups** | **% proteins in orthogroups** |
| --- | --- | --- | --- |
| *Acyrthosiphon pisum* | 34843 | 32076 | 92.1 |
| *Aedes aegypti* | 40355 | 38508 | 95.4 |
| *Callosobruchus maculatus* | 30616 | 27114 | 88.6 |
| *Cimex hemipterus* | 16500 | 14341 | 86.9 |
| *Cimex lectularius* | 23323 | 22487 | 96.4 |
| *Drosophila melanogaster* | 22117 | 19880 | 89.9 |
| *Gryllus bimaculatus* | 25719 | 19283 | 75 |
| *Rhodnius prolixus* | 30931 | 28467 | 92 |
| *Triatoma infestans* | 18543 | 16519 | 89.1 |
| Dmel Wolbachia | 1172 | 991 | 84.6 |
| Clec Wolbachia | 1061 | 947 | 89.3 |

***Table S2.*** *Reference table for characterising Sperm-leucylaminopeptidase (S-Lap) catalytic window motif.*

| **blLAP role** | **S-Lap position** | **Script site** | **Offset** | **Compatible** | **Evidence [reference]** |
| --- | --- | --- | --- | --- | --- |
| Lys250: Site 1 Zn coordination via amino group | Lys 327 | Metal 1 | 0 | Gln | Gln observed in active S-Laps (Dorus et al. 2011); amide N substitutes for amino N |
| Asp255: Site 2 tight Zn coordination via carboxylate | Asp 332 | Metal 2 | 5 | Asn, Glu, His, Cys | Asn/Glu conservative oxygen donors; His coordinates via imidazole N (loopin, (Dorus et al. 2011)); Cys thiol coordinates (S-Lap5, (Dorus et al. 2011)) |
| Asp273: Site 1 loose Zn coordination via carboxylate | Asp 350 | Metal 3 | 23 | Asn, Glu, Ser, Thr, Cys | Asn/Glu conservative oxygen donors; Ser/Thr hydroxyl O can coordinate divalent metals; Cys thiol coordinates |
| Lys 409: Site 1 loose Zn coordination via amino group | Lys 409 | Metal 4 | 82 | Asp, Ser | Lys -> Asp and Lys -> Ser observed in active S-Laps (Dorus et al. 2011); less mutagenesis constrained than Glu334 |
| Glu344: Bridges both Zn sites via two carboxylate oxygens | Glu 411 | Metal 5 | 84 | Asp | E -> D retains >95% activity ((Gu and Walling 2002) tLAP-A Glu 429); E->V and E->S retain <3% - not compatible |
| Lys262: Catalytic base; activates nucleophilic water | Lys 339 | Cat 1 | 12 | - | Most substitutions reduce activity (Gu and Walling 2002) |
| Arg336: Transition state stabilisation of gem-diolate | Arg 413 | Cat 2 | 86 | - | Essential for catalysis (Straeter and Lipscomb 1995) |

**blLAP role**: residue (position in mature protein sequence) for M17 leucyl-aminopeptidase reference sequence derived from blLAP crystal structures (PDB 1LAM (Burley et al. 1992)).

**S-Lap position**: corresponding position for S-Laps for *D. melanogaster* (Dorus et al. 2011).

**Offset**: number of residues upstream from the start of motif (Lys 327).

**Compatible**: other compatible residues at the given site.

**SUPPLEMENTARY RESULTS**

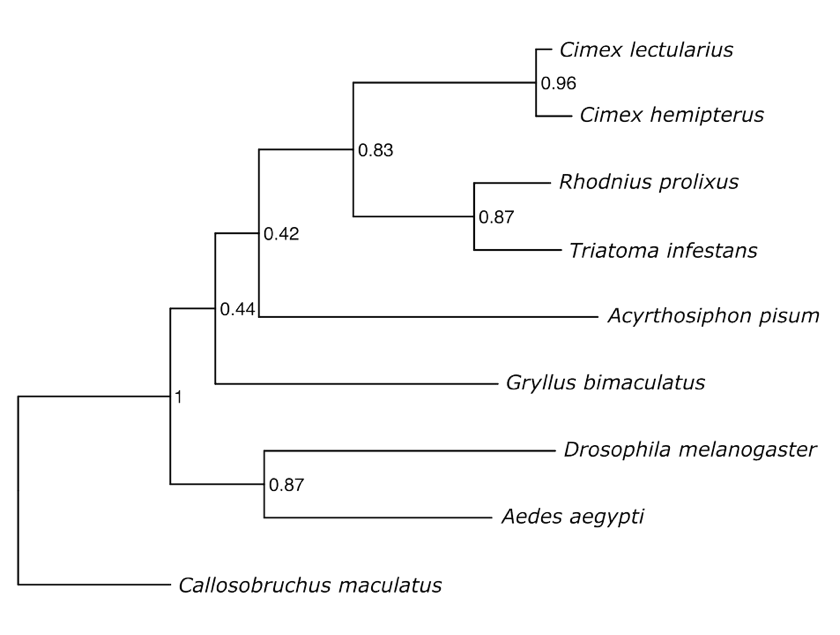

***Figure S1.*** *Phylogeny of the species using for orthologue comparisons from OrthoFinder rooted using STRIDE (Emms and Kelly 2015; Emms and Kelly 2019). Node labels show STAG support values.*

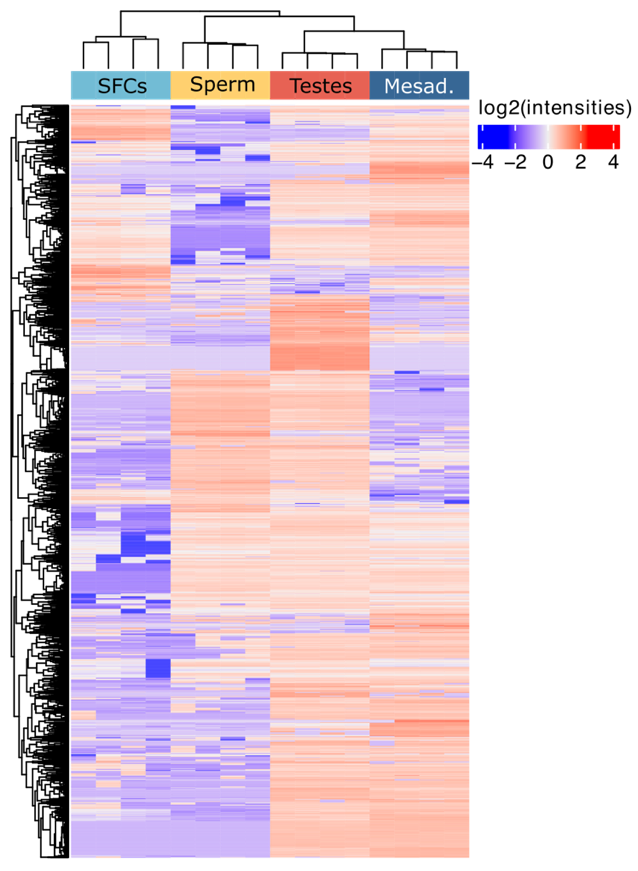

***Figure S2.*** *Heatmap showing z-scored log2(abundance) for proteins included after filtering (n = 3348) for each tissue/sperm and replicate (n = 4).*

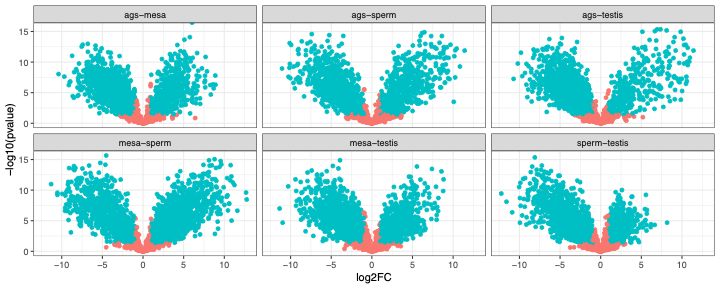

***Figure S3.*** *Volcano plots showing pairwise differences between each tissue. Tissue bias was defined based on a log2FC > 1 and p < 0.05 (blue) in that tissue compared to all others.*

***Endosymbiont/bacteriome proteins.*** *Wolbachia* and other endosymbiotic bacteria (e.g., *Rickettsia*) (Thongprem et al. 2020) are localised in bacteriomes (aka mycetomes) attached to the *vas deferens* at the base of the testes (Usinger 1966; Hosokawa et al. 2010). We identified 176 endosymbiont proteins (149 *Wolbachia*, 27 *Rickettsia*) in the full dataset (n = 4651). Applying less stringent filtering criteria than above (≥ 1 unique peptide) retained 110 *Wolbachia* proteins and 22 *Rickettsia* proteins. As expected, the majority of *Wolbachia* proteins showed highest abundance in the testes/bacteriome samples (Fig. S4). The subset of *Wolbachia* proteins we identified (110/1027; 10.7%) showed enrichment for GO terms including: translation (*p* < 0.001), regulation of gene expression (*p* = 0.016), proteolysis (*p* = 0.016) (BP); cytosolic ribosome (*p* = 0.006), large ribosomal subunit (*p* = 0.007), ribonucleoprotein complex (*p* = 0.007) (CC); structural constituent of ribosome (*p* < 0.001), protein binding (*p* = 0.005), RNA binding (*p* = 0.006) and rRNA binding (*p* = 0.012) (MF). All but three (107/110; 97.27%) *Wolbachia* proteins in *C. lectularius* that we identified had orthologs in *Wolbachia* in *Drosophila, as* identified using *OrthoFinder*.

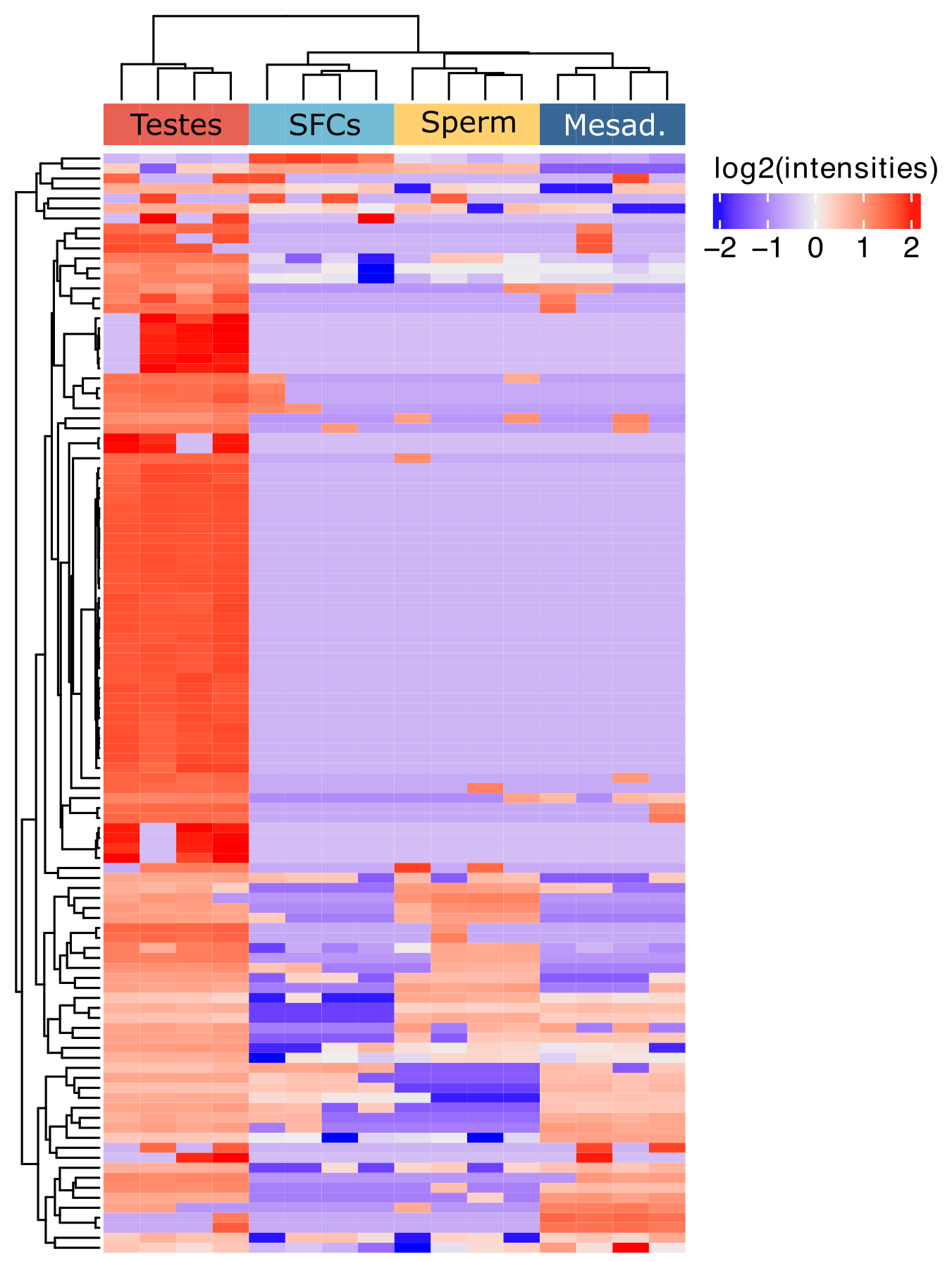

***Figure S4.*** *Abundance (z-transformed log2-normalised abundance) of Wolbachia endosymbiont proteins (n = 110) in male reproductive tissues.*

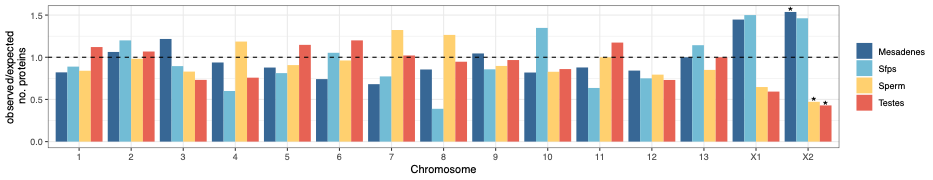

***Figure S5.*** *Chromosomal distribution of male-biased proteins on the autosomes and each of the two* C. lectularius *X chromosomes. Asterisks represent results from comparisons of the observed to the expected number of proteins on each chromosome after multiple testing correction using* $X^{2}$ *tests. The dashed line indicates the null expectation.*

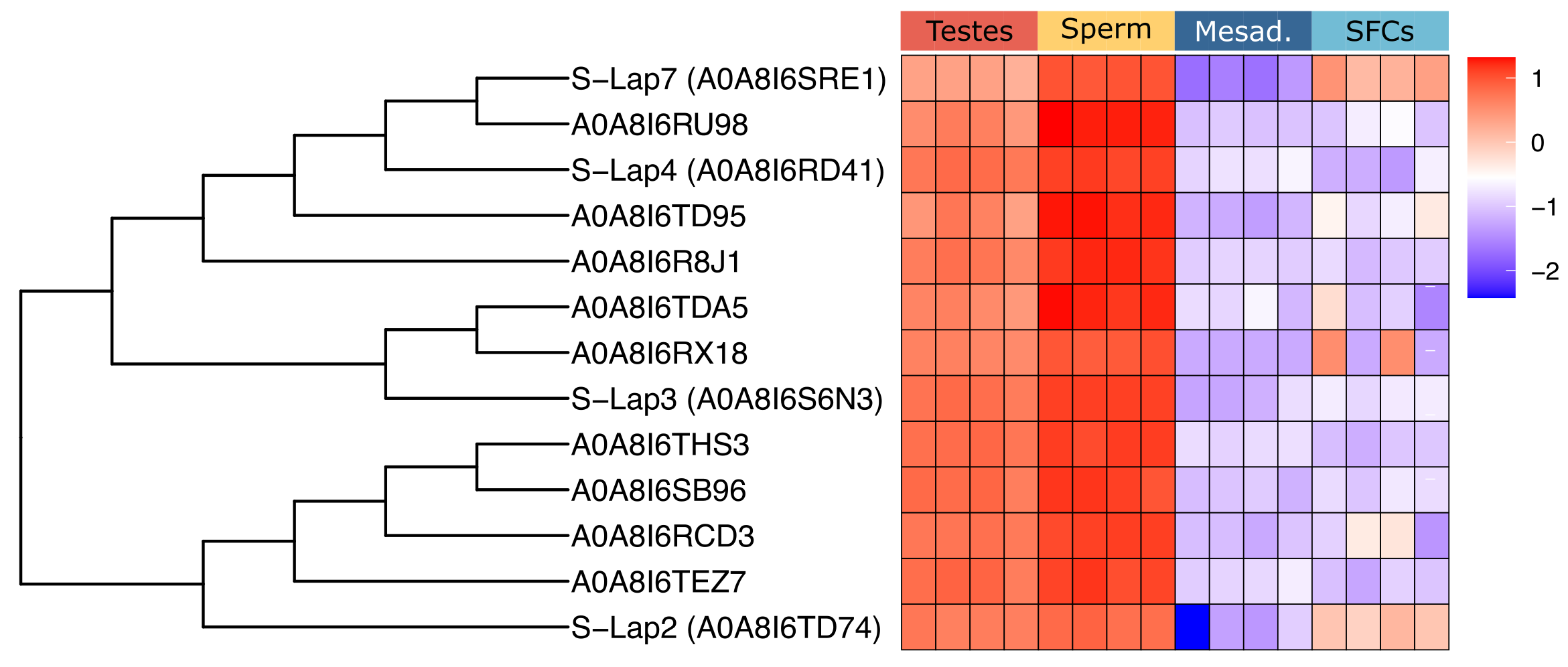

***Figure S6.*** *z-scored log2(abundance) of* C. lectularius *S-Lap orthologs in male reproductive tract samples. Gene tree on the left-hand side shows a pruned gene tree for* Cimex *S-Laps with one-to-one orthologs with* D. melanogaster *labelled.*

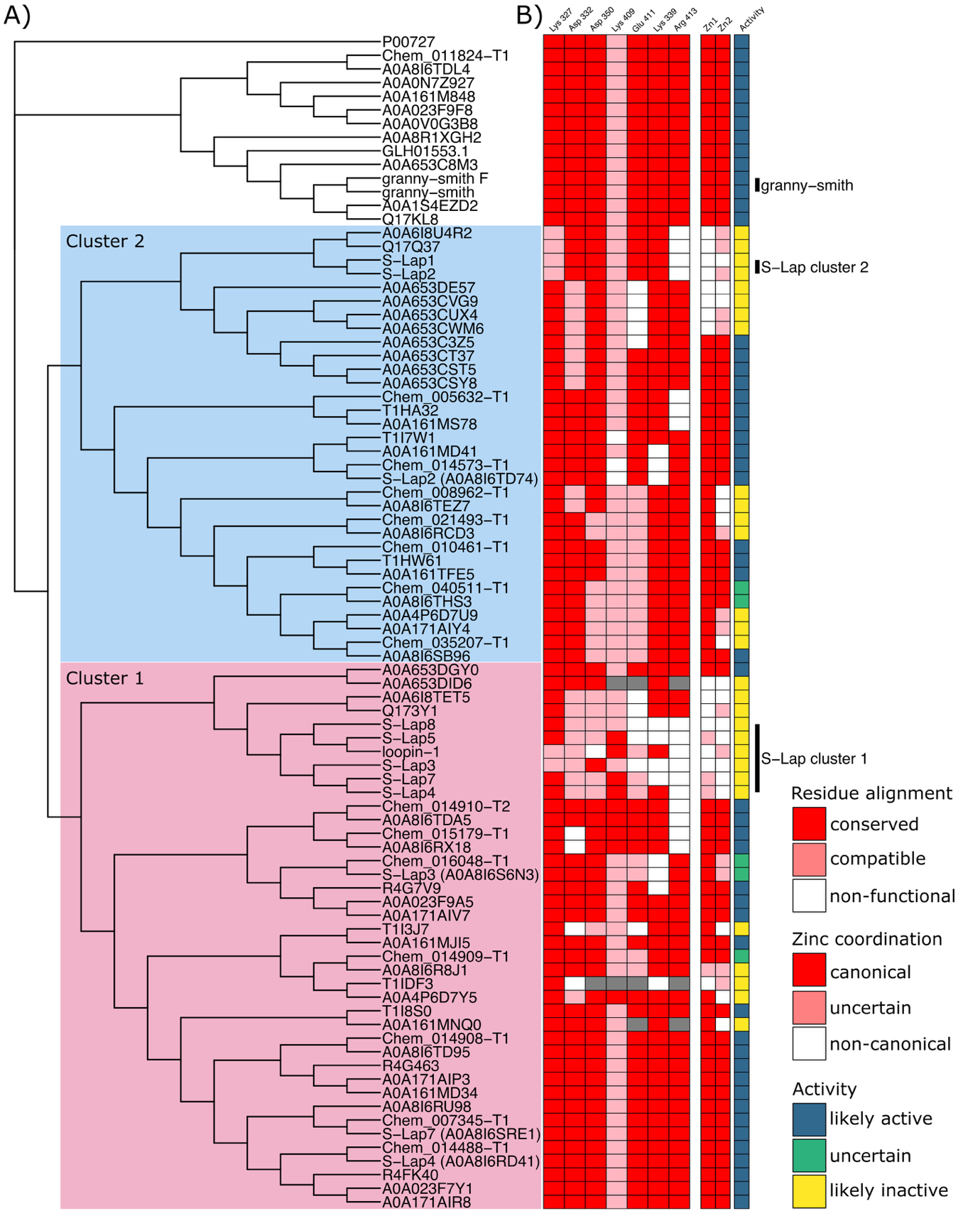

***Figure S7.*** *Catalytic activity in S-Lap and granny-smith orthologs.* ***A)*** *Gene tree for S-Lap and granny-smith orthologs identified using OrthoFinder. Protein sequences were aligned using MAFFT (Madeira et al. 2024) and bootstrapped 1,000 times.* ***B)*** *Alignment of S-Lap and granny-smith orthologs focused on amino acid residues against the M17 amino peptidase metal binding motif. Amino acids are coloured by alignment to consensus (conserved, red; compatible, pink; non-functional, white). Zn^2+^ ion binding at the “loose” binding site (Zn1) and “tight” binding site (Zn2) is scored as canonical (red), uncertain (pink) or non-canonical (white). Overall activity is scored as likely active (blue), uncertain (green), or likely inactive (yellow) based on coordinating residue geometry, number and inter-ligand angles and inter-zinc distances measured from AlphaFold3 models in ChimeraX.*

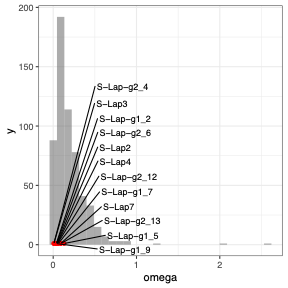

***Figure S8.*** *Rates of molecular evolution (pairwise dN/dS between C. lectularius and C. hemipterus) for S-Lap orthologs. Histogram shows dN/dS values for sperm proteins (n = 580) and red points show S-Lap orthologs labelled based on one-to-one orthology with* D. melanogaster *or membership to S-Lap cluster 1 (g1) or 2 (g2)* (Dorus et al. 2011) *followed by a unique ID.*
